# Potentials of Machine Learning in Predicting Key Features of Synthetic Antimicrobial Polymers (SAMPs)

**DOI:** 10.1101/2025.08.24.671690

**Authors:** Lena Dalal, Deborah Stolte Bezerra Lisboa De Oliveira, Nicholas J. Warren, Olivier Cayre, Sebastien Perrier

## Abstract

The widespread antimicrobial resistance urged the need for novel antimicrobial agents. Synthetic antimicrobial polymers (SAMPs) were proposed as promising antibiotics to overcome the drawbacks of host-defence peptides. A machine learning forecast can be beneficial to evaluate the influence of SAMPs features on their potency and toxicity. In this study, we utilised a library of 20 polyacrylamides varied in: 1) type of amine side chain, 2) chain length, 3) cationic amine ratio, and 4) polymer architecture. Their structure-activity relationship was evaluated by comparing experimental observations and machine learning models. While classification models showed good fit in the training set, regression models demonstrated better fit in the testing set. Regression random forest and gradient boosting methods demonstrated reliable reproducibility of feature importance and maintained the tree structure throughout multiple runs. Beeswarm and waterfall plots provided an overview of the joint SHapley Additive exPlanations (SHAP) values of features on a specific data point. Based on validation tests and the consistency of feature importance and SHAP values, we conclude that boosting-ensemble methods can be utilized in forecasting future SAMPs.

## Introduction

Antimicrobial resistance has become a significant global concern, and the development of new generations of antibiotic is crucial for the coming future.^1^ This pressing issue has piqued the interest of many researchers during the last decade.^2^ Since the discovery of host-defence peptides (HDPs) in the 1980s, they have been considered a promising broad-spectrum antibiotics.^3^ HDPs are short peptides, consisting of a combination of less than 50 amino acids in secondary structure conformation. Their amino acid combination is crucial to their antimicrobial and immune response, as well as their selectivity towards microbial cells.^4^ HDPs comprise of a mixture of hydrophobic and cationic amino acid distributed to yield a net positive charge at physiological pH.^5^ This unique amphiphilic structure enables them to be soluble in the physiological aqueous environment, while maintaining their ability to partition within phospholipid microbial membrane.^6^ Moreover, the dominant cationic charge of HDPs prompts electrostatic interactions with the microbial cell membrane dominated by negatively charged lipids. These interactions lead to disruption of the membrane structure, formation of pores and eventually apoptosis.^7,8^ However, mammalian cell membrane consists of zwitterionic phospholipids (i.e., phosphatidylglycerol, cardiolipin) oriented outward, which minimises interactions with HDPs, thus promoting the peptides’ selectivity towards microbial cells.^9–11^

This has prompted an extensive search for synthetic antimicrobial peptides mimics (SAMPs) that replicate the activity of HDPs, while offering enhanced stability and tuneable chemical structure. Among the most versatile SAMPs reported to date are polymers functionalized with amino acid-mimicking groups,^12,13^ built on diverse backbones including (but not limited to) polynorbornene^14^, polyethyleneimines^15^, polynylon-3^16–18^, polyvinylpyrrolidones^19^, Poly(2-oxazoline)s^20,21^ polyurea^22^, methacrylates^23–27^ and acrylamides^27–29^.

Many structural parameters were found to affect SAMPs potency, in particular molecular weight, polymer backbone structure, type of cationic moieties and its ratio to the hydrophobic monomer of choice, as described by Hartlieb *et al.*^30^, Pham *et al.*^31^ and Locock *et al*.^24,32^ who focused mainly on copolymers of methacrylate and acrylamides derivatives. Well-defined antimicrobial polymethacrylates or polyacrylamides can be obtained via controlled radical polymerization techniques such as copper(0)-mediated radical polymerisation^33^ and reversible addition-fragmentation chain transfer polymerization (RAFT).^34–36^ RAFT polymerisation not only provides a good control over the polymers molecular weight but also allows for telescoping synthesis of multiblock with narrow dispersity.^37^

In previous work by Perrier group^38,39,40,41,42^ polyacrylamides selected to resemble the amino acids functional groups found in innate HDPs (e.g. lysin, arginine and isoleucine) showed great potential for future topical antimicrobials mimicking HDPs with improved resilience. These monomers’ high propagation rates (kp) facilitate their rapid and controlled polymerization via RAFT polymerisation, allowing for the control of the cationic moieties distribution along the polymeric chain, to achieve the desired amphiphilic balance of HDPs with higher stability against degradation.^43^

In our recent study^44^, we concluded the main factors influencing the therapeutic profile of acrylamide-derived SAMPs include the type of cationic moieties and its ratio, the polymer chain length and sequence. However, elucidating structure-activity relationship (SAR) and optimising the design of antimicrobial polymers is a time-consuming process, which calls for a the utilisation of automated methods.^24,30–32^

The recent advances in artificial intelligence tools provide a remarkable path towards more efficient and robust SAR analysis and materials design. Particularly, machine learning (ML), a subset of AI which involves algorithms designed to generate predicted outcomes from historic data, appears to be a powerful tool to guide the design of the next generation of antimicrobials.^45,46^ In this approach, *in silico* screening minimizes the load of experimental work required, as it can narrow down the number of potential candidates by predicting activity patterns.^47^ It can also foresee hidden patterns when screening drug candidates database, e.g. antibiotics, that otherwise might be missed as prospective candidates.^48^ The application of ML in the high-throughput screening of antibiotics candidates libraries has been reported recently^49–53^, but a number of challenges are still to be addressed.^54^ The typical workflow for ML includes data collection, features engineering, model selection and validation, and finally model application.^55^ Data can be sourced from direct experimental work, published literature, or existing databases. Experimental data have the advantage of controlled parameters, particularly for polymers, ensuring consistency and reliability. However, it is time-consuming and typically yields smaller datasets. In contrast, utilizing published research provides access to extensive datasets within a shorter timeframe, but the diversity of experimental methods and data reporting standards may impact the quality and accuracy of ML predictions.^55,56^ Moreover, the accuracy of predictions rely on identifying relevant input variables (that is, design features) that influence output performance. Hence, it is critical to consider the relevant features or descriptors to be utilised in the ML pipeline in an early stage, based on experimental knowledge but preferably prior to extensive data collection. This raised a challenge for HDPs, when screening through their wide range of structures and sequences,^54,57^ and similarly, it could be limiting for the use of ML models for SAMPs as well.

Another challenge is selecting an appropriate ML model, as identifying the optimum model for the dataset is critical for successful predictions. There are many ML techniques to choose from for medicinal chemistry, i.e. decision tree, random forest, gradient boosting, and artificial neural networks, including algorithms for both regression and classification methods.^48^ Model selection considers the nature of the dataset itself, computational time restrictions, and additional insights that may be available for each specific model. One insight which can be obtained from ML models is feature importance, which gives the contribution of each feature to the model output and can facilitate the identification of SAR. Permutation feature importance can be performed after model selection and fitting and can be applied to any ML model. However, this requires additional calculations and consumes more computational time. Tree-based models, on the other hand, calculate feature importance innately as part of their fitting algorithm, reducing the computational effort needed.^58^

Herein, we have utilised decision tree models and two tree-ensemble methods: random forests (RF) and gradient-boosting (GB). Tree-based models were selected as there is no additional computational expense in calculating feature importance for these models. Decision trees are branching structures that can be used for prediction based on splitting point (node) rules. In its simplest form, a single decision tree is the predictor. The structure of the tree (node feature selection and threshold value) is tuned by using known data, and either minimizing the GINI impurity (for classification) or the “within-sample” variance (for regression method) at each node.^59,60^ Equations describing how this structure tuning occurs are shown in the methods section.

RF are an ensemble ML technique that combines the prediction of different decision trees, thus minimizing overfitting and data bias.^61^ This combination is also referred to as bagging, and considers the predictions of all trees in the forest, either by averaging the final value (for regression) or by majority counting of the selected class (for classification).

GB is another type of tree ensemble method. Unlike bagging, boosting considers trees in sequence, with each new tree building upon the previous one to improve the fit. During tree structure creation and tuning, GB uses the error between expected and calculated output values, instead of the output value itself. The new predicted output value can then be determined from the previous prediction and the new predicted error, until the prediction does not change. The mathematical development is explained in the work published by Friedman.^62^

After applying the appropriate ML model to the dataset, analysing the contributions of each input feature can be done via feature importance (FI) calculations in tree models and SHapley Additive exPlanations (SHAP) analysis.^63^ FI gives the overall contribution of each feature to the output. However, FI does not provide any information on whether the contribution is negative or positive, or on what each features’ contribution is for specific datapoints. Additional information can be gained by using SHAP values, which give the marginal contribution of each feature for each datapoint in the dataset, allowing for the identification of features and feature values that lead to higher-performing SAMPs designs, as well as clustering of SAMPs designs that have similar feature contribution distributions.^63^

Herein, we ran a pilot study, where a ML pipeline was applied to a small set of controlled experimental data with 4 variables. The work includes the use of 3 algorithms: decision tree and its ensemble methods, random forest and gradient boosting, applied as both regression and classification methods. We explored their potentials for the prediction of antimicrobial properties of a library of polyacrylamides.

All the polymers in the experimental dataset are polyacrylamides with the same chain transfer agent (CTA), and *N*-Isopropyl acrylamide (NIPAM) as the apolar monomer of choice, to facilitate comparison. The studied SAMPs were designed to vary in degree of polymerization, architecture, type of cationic moiety, and cationic ratio, to enable a broad range of experimental data to be used with the chosen algorithms.

The output variables of the pipeline were 1) the minimum inhibitory concentration (MIC) and 2) hemagglutination, used to predict antimicrobial activity and identify the most significant feature for the optimal selectivity, respectively. As structure-activity relationship was also related to the type of bacteria tested, the minimum inhibitory concentration (MIC) of each tested strain was used as an output.

In this work we trained the algorithms and identified the model with the best performance, which provides reproducible feature importance results after multiple runs. The aim is to advance our understanding of structure-activity relationships of antimicrobial polyacrylamide without the need for further experimental work. To the best of our knowledge, such study has only been done recently by Kundi *et al.*^64^ Their work used a previously published dataset and utilised a decision tree classification model using multiple descriptors, and in conclusion provided recommendations for future design of SAMPs.^64^

## Experimental

### Polymers synthesis and bioassays

The polymers used in this work were made via RAFT polymerisation. Their synthesis process and characterisation as well as the minimum inhibitory concentration (MICs) against Gram-negative *P. aeruginosa* (PA14 and LESB58) and Gram-positive *S .aureus* (USA300 and Newman) strains, and hemocompatibility (C_H_) were all detailed in a previously published study.^44^

### Machine learning techniques

#### Mapping input-output relationships with random forest feature importance and SHAP values

Machine learning tools were applied to the data to relate the contribution of each input variable (feature) to each output variable. Separate models were generated for each output (MIC PA14, MIC LESB58, MIC USA300, MIC Newman and agglutination) based on 4 inputs (type and percentage of cationic monomer, degree of polymerization, and polymer conformation). Three methods were used to create these models: single decision trees, random forests (RF) and gradient-boosting (GB). The complete dataset for model fitting and validation is presented in Table 1 and Table 2. It should be noted that categorical inputs (type of cationic monomer and polymer conformation) were assigned a numerical value, and that, while experimental data is available for hemolysis, it was not possible to create models for this output variable because all output values were the same (>512 µg/ml).

**Table 1.** Dataset for all four MIC output variables (µg/mL)

| Data point # | DP | CatMonPerc | PolyConf* | CatMonType** | MIC PA14 | MIC LESB58 | MIC USA300 | MIC Newman |
| --- | --- | --- | --- | --- | --- | --- | --- | --- |
| 1 | 25 | 70 | 2 | 1 | 513 | 128 | 128 | 128 |
| 2 | 25 | 30 | 2 | 1 | 513 | 512 | 513 | 513 |
| 3 | 25 | 30 | 2 | 2 | 513 | 513 | 513 | 513 |
| 4 | 25 | 30 | 2 | 3 | 513 | 513 | 513 | 513 |
| 5 | 50 | 100 | 1 | 2 | 513 | 513 | 64 | 64 |
| 6 | 50 | 100 | 1 | 3 | 513 | 513 | 128 | 256 |
| 7 | 50 | 100 | 2 | 3 | 513 | 513 | 128 | 256 |
| 8 | 50 | 70 | 2 | 1 | 128 | 64 | 64 | 64 |
| 9 | 50 | 70 | 2 | 2 | 513 | 513 | 513 | 513 |
| 10 | 50 | 70 | 2 | 3 | 513 | 513 | 513 | 513 |
| 11 | 50 | 50 | 2 | 1 | 512 | 512 | 256 | 256 |
| 12 | 50 | 50 | 4 | 1 | 513 | 513 | 513 | 513 |
| 13 | 50 | 50 | 2 | 2 | 513 | 513 | 513 | 513 |
| 14 | 50 | 50 | 2 | 3 | 513 | 513 | 513 | 513 |
| 15 | 50 | 30 | 3 | 1 | 64 | 256 | 513 | 513 |
| 16 | 50 | 30 | 2 | 1 | 513 | 512 | 513 | 513 |
| 17 | 50 | 30 | 4 | 1 | 513 | 513 | 513 | 513 |
| 18 | 50 | 30 | 3 | 2 | 513 | 513 | 513 | 513 |
| 19 | 50 | 30 | 2 | 2 | 513 | 513 | 513 | 513 |
| 20 | 50 | 30 | 3 | 3 | 513 | 513 | 513 | 513 |
| 21 | 50 | 30 | 2 | 3 | 513 | 513 | 513 | 513 |
| 22 | 100 | 30 | 2 | 1 | 256 | 128 | 128 | 128 |
| 23 | 100 | 30 | 4 | 1 | 256 | 128 | 513 | 513 |

\***Polymer conformation:** 1 = Homopolymer, 2 = Diblock copolymer, 3 = Triblock copolymer, 4 = statistical copolymer. \*\***Type of cationic monomer:** 1 = AEAM, 2 = DMAEAM, 3 = TMAEAM

**Table 2.** Dataset for agglutination (µg/mL)

| Data point # | DP | CatMonPerc | PolyConf* | CatMonType** | Agglut |
| --- | --- | --- | --- | --- | --- |
| 1' | 25 | 70 | 2 | 1 | 4 |
| 2' | 25 | 30 | 2 | 1 | 32 |
| 3' | 25 | 30 | 2 | 2 | 256 |
| 4' | 25 | 30 | 2 | 3 | 16 |
| 5' | 50 | 100 | 2 | 3 | 64 |
| 6' | 50 | 70 | 2 | 1 | 4 |
| 7' | 50 | 70 | 2 | 2 | 513 |
| 8' | 50 | 50 | 2 | 1 | 513 |
| 9' | 50 | 50 | 4 | 1 | 128 |
| 10' | 50 | 50 | 2 | 2 | 513 |
| 11' | 50 | 30 | 3 | 1 | 513 |
| 12' | 50 | 30 | 2 | 1 | 32 |
| 13' | 50 | 30 | 4 | 1 | 128 |
| 14' | 50 | 30 | 3 | 2 | 513 |
| 15' | 50 | 30 | 2 | 2 | 256 |
| 16' | 50 | 30 | 3 | 3 | 256 |
| 17' | 50 | 30 | 2 | 3 | 128 |

\***Polymer conformation:** 1 = Homopolymer, 2 = Diblock copolymer, 3 = Triblock copolymer, 4 = statistical copolymer. \*\***Type of cationic monomer:** 1 = AEAM, 2 = DMAEAM, 3 = TMAEAM

All three methods were tested as classification and regression methods, that is, treating the output variables as categorical or numerical, respectively. Previous works that modelled polymer property-performance with ML have used classification models.^64–66^ However, for the dataset in the present work, it was found that classification methods were not reproducible within different runs with distinct training-validation dataset splits, due to uneven class distribution in the dataset (see Results section and Supplementary information for further discussion). Therefore, regression methods were tested as well. For classification methods, continuous output values were assigned either to class 1 (good performance) or class 0 (undesirable performance), with a continuous output value of 64 µg/ml used as the threshold value. Depending on the output variable, good performance is attributed to values ≤ 64 µg/ml, or > 64 µg/ml. For example, for potency, lower concentrations are desired, hence for all MIC outputs, values ≤ 64 µg/ml were assigned to class 1. Conversely, it is undesirable to induce hemagglutination at lower concentrations, so values > 64 µg/ml were assigned to class 1 in this case.

### Machine learning models

For regression decision tree fitting, 20% of the dataset was used for testing/validation. For classification tree fitting, the training/testing split was 50% to ensure both classes was present in the training dataset. This was performed in Python with the scikit-learn package.^58^

The scikit-learn package for Python was used to create RF ensembles, with out-of-bag cross-validation and hyperparameter optimisation.^58^ 20% of the datapoints were randomly selected with replacement and used to validate each tree structure in the forest for the regression method. As the dataset is small, cross-validation was chosen instead of splitting between training-test datasets for regression. The split was 50% for classification. Fitting was repeated 5 times to ensure reproducibility.

Similarly to the other two methods, GB was implemented in Python with scikit-learn, using 20% of the dataset for cross-validation with the regression method, and a 50% train-test split for classification.

### Feature importance and SHAP values

SHAP values calculations were implemented in Python with the SHAP package.^67^ SHAP was used to explain how each feature contributed to the predicted output values of the best previously generated models. More information on how feature importance is calculated for tree models is given below.

### Tree models and feature importance

Figure 1 shows a split across a node t in a decision tree T, for continuous output data. When fitting the tree structure, the goal is to minimise the impurity across the node. This is accomplished by selecting which feature the node is splitting on (X_n_) and what threshold value is used for the split condition (s).

**Figure 1.**
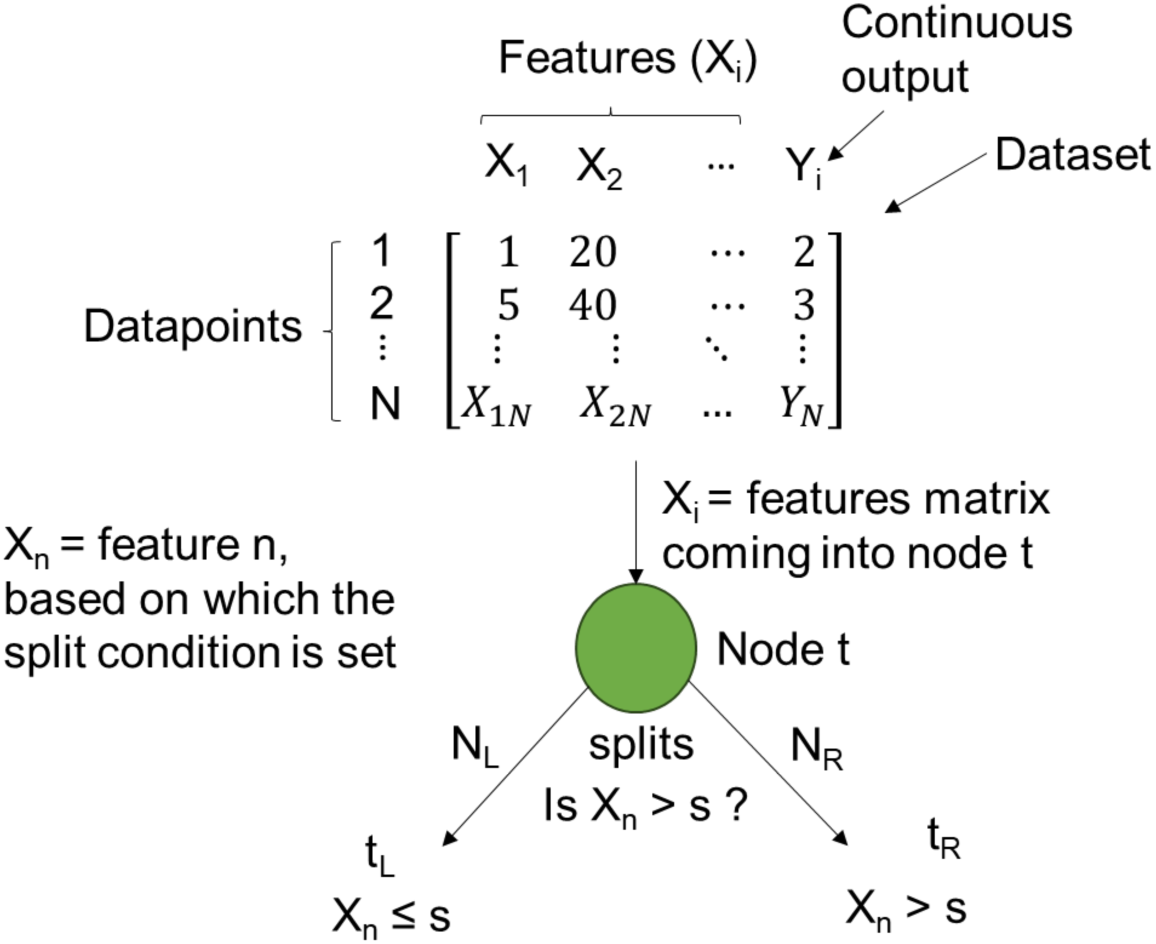
Sample node t within a tree T. Coming into node t is matrix X_i_, containing information about the features (input variables). If node t is the first node in the tree, then this matrix contains all N datapoints in the dataset. Otherwise, matrix X_i_ contains a subset that comes from a previous node split. X_n_ is the feature over which the dataset splits, and s is the threshold value for this split. A subset of matrix X_i_ (with N_L_ datapoints) splits to the left at node t split t_L_, for the datapoints in which feature X_n_ has a value equal to or smaller than the threshold s. Another subset of X_i_ (with N_R_ datapoints) splits at t_R_, for values of X_n_ greater than the threshold. Within a tree T, the selection of X_n_ and s for each node split is achieved by minimising the impurity of the node with respect to the output variable, Y_i_.

The concept of impurity comes from classification trees, where impurity represents the degree to which datapoints have been incorrectly classified at any node. For regression trees, impurity is treated as the variance, or “within-sample” variance^68^, as shown in the equations below.

The impurity 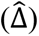 coming into node t is given by:

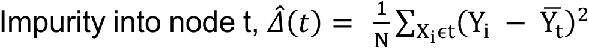

Where N is the number of samples (datapoints) in node t; Y_i_ is each output value present in the dataset coming into node t; and 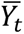 is the sample mean for the output variable for datapoints coming into node t.

The impurity coming out of node t is:

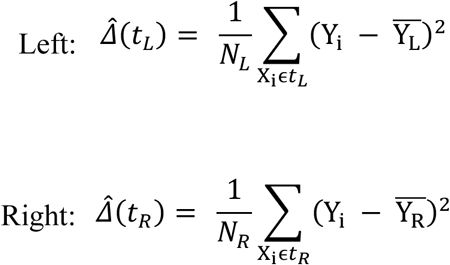

The variables are similar to the ones coming into the node, except now N_L_ and N_R_ are the number of datapoints present in the left and right splits of the node, respectively; Y_i_ is each output value in the part of the incoming dataset that was split into either the left or right split; and 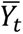 and 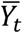 are the mean output values for datapoints in the left and right splits, respectively.

The decrease in impurity across node t, is given by the difference in impurity coming in and out, with the impurity coming out weighted by the fraction of samples that go into each split.

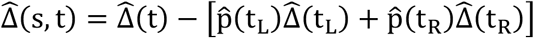

Where:

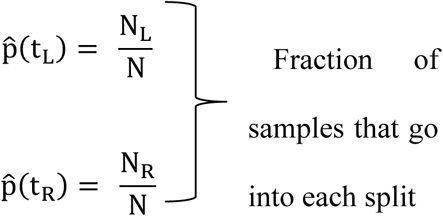

To minimise impurity, the algorithm maximises 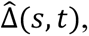 which is the *decrease* in impurity.

For classification trees, “within-sample” variance is replaced by GINI impurity (given below) in the equations above.

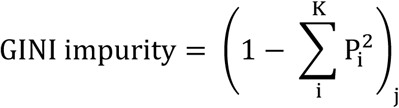

Where K is the total number of classes present (usually two for a binary classification problem), i is each class, and P_i_ is the probability of observing class i in split j.

Feature importance for each node t (I_node,t_) can be calculated directly from each splits’ impurity.

This is shown below for a classification tree.^69^

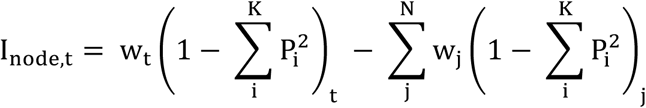

Where t is the node; w_t_ is the weight of the sub-dataset coming into node t (that is, the fraction of the total datapoints in the dataset that belong to this sub-dataset); subscript t represents the sub-dataset coming into the node; w_j_ is the weight of the splits coming out of node t (for the usual case where each node is a bifurcation, N=2, and j = 1 or 2, left or right); and subscript j represents each split out of node t.

The importance of each feature n can then be calculated by summing the importance of all nodes which split based on feature n, and then dividing by the sum of the importance of all nodes.

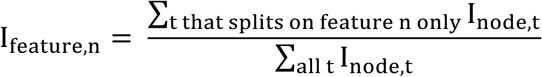

Feature importance is then normalised between 0 and 1.

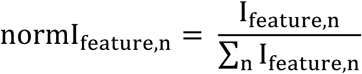

For bagging ensemble methods, such as random forests, the combined feature importance over all trees is given by ^58^:

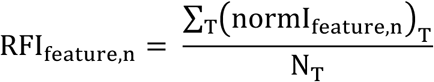

Where T is each tree in the forest, and N_T_ is the total number of trees.

### Model fit and consistency testing

To select the best performing model(s), all three methods (decision tree, RF and GB) as both regression and classification, were fit in 5 different runs (i.e., with different random selections for training and testing datasets) while optimizing some of each method’s hyperparameters. Hyperparameters that affect the size of the trees and forests were chosen for optimization (such as maximum tree depth and number of estimators). Hyperparameters related to the mathematical formulation of the algorithm were left at their default values and those related to the selections made in regard to testing and validation splits, cross-validation, and bagging were set to their specific values as mentioned previously in this section. It should be noted that tuning different hyperparameters can lead to similar model results, and this case, the hyperparameter tuning that gives the smaller training error was chosen.^58^

The fits of each method (expressed as the R^2^ value for regression, and precision, recall and F1 score metrics for classification) were considered when selecting which method would be best for representing the polymer property-performance space. However, as it was noticed that some methods exhibited different feature importance distributions among different runs, or different tree structures for single decision tree and the final gradient boosting tree, the ability of each method to remain consistent across different runs was also factored into the model selection decision.

## Results

### Polymer library

A total of 20 polymers (Table 3) were utilised with NIPAM as the apolar monomer of choice for all copolymers and the cationic monomers based on 1) the primary amine ethyl acrylamide (AEAM), 2) the tertiary amine dimethyl ethyl acrylamide (DMAEAM) and 3) the quaternary amine trimethyl ethyl acrylamide (TMAEAM). The cationic monomer ratios were 30, 50, 70, and 100%. The polymers characterisation results including ^1^HNMR and GPC and the minimum inhibitory concentrations (MICs) against *P. aeruginosa* and *S. aureus strains*, and hemagglutination (C_H_) concentrations were all determined in previously published work.^44^

**Table 3.** The library of polymers used.

| **P** | **CatMon\*** | **DP<sub>target</sub>** | **Polymer Architecture** | **$f_{\text{CatMon}}$** |
| --- | --- | --- | --- | --- |
| **P1** | AEAM | 25 | Diblock | 0.3 |
| **P2** | AEAM | 25 | Diblock | 0.7 |
| **P3** | AEAM | 50 | Statistical | 0.3 |
| **P4** | AEAM | 50 | Statistical | 0.5 |
| **P5** | AEAM | 50 | Diblock | 0.3 |
| **P6** | AEAM | 50 | Diblock | 0.5 |
| **P7** | AEAM | 50 | Diblock | 0.7 |
| **P8** | AEAM | 50 | Triblock | 0.3 |
| **P9** | DMAEAM | 25 | Diblock | 0.3 |
| **P10** | DMAEAM | 50 | Diblock | 0.3 |
| **P11** | DMAEAM | 50 | Diblock | 0.5 |
| **P12** | DMAEAM | 50 | Diblock | 0.7 |
| **P13** | DMAEAM | 50 | Triblock | 0.3 |
| **P14** | TMAEAM | 25 | Diblock | 0.3 |
| **P15** | TMAEAM | 50 | Diblock | 0.3 |
| **P16** | TMAEAM | 50 | Diblock | 0.5 |
| **P17** | TMAEAM | 50 | Diblock | 0.7 |
| **P18** | TMAEAM | 50 | Triblock | 0.3 |
| **P19** | DMAEAM | 50 | Homo | 1 |
| **P20** | TMAEAM | 50 | Homo | 1 |

**CatMon:** the cationic monomer. **NIPAM:** *N*-isopropyl acrylamide. **AEAM:** *N*-(2-aminoethyl) acrylamide. **DMAEAM:** *N*-[2-(dimethylamino)ethyl] acrylamide. **TMAEAM:** *N*-[2-(trimethylamino)ethyl] acrylamide. **DP:** degree of polymerisation. **$f_{\text{CatMon}}$ :** the fraction of cationic monomer to the apolar

Briefly, the results highlight the significance of cationic charge in determining the activity and selectivity of polymers. Using the primary amine in high ratio (70%) in diblock copolymer enhanced the antimicrobial profile but reduced the selectivity. Polymer architecture also influenced the potency, AEAM-triblock (DP50-Ta30) induced similar potency towards Gram-negative bacteria strains at lower cationic ratio (30%) than the diblock (DP50-Da70). The methylation of the cationic units of the block copolymers diminished their potency. Only homopolymers of DMAEAM and TMAEAM were active against Gram-positive bacteria.

### Machine learning model selection

Model selection considered both the fit and reproducibility of feature importance.

### The fit and predictability of the ML methods

For each method used to create models (decision tree, RF, and GB), the fit was evaluated as the R^2^ value for regression methods (Table 4), and precision, recall and F1 scores for classification methods (Table 5).

**Table 4.** Fit metrics of the regression models.

| Method | Output variable | R<sup>2</sup>* |
| --- | --- | --- |
| Decision tree | MIC PA14 | 1.00<sup>+</sup> |
|  | MIC LESB58 | 1.00<sup>+</sup> |
|  | MIC USA300 | 1.00<sup>+</sup> |
|  | MIC Newman | 1.00<sup>+</sup> |
|  | Agglutination | 1.00<sup>+</sup> |
| Random forest | MIC PA14 | 0.82 |
|  | MIC LESB58 | 0.90 |
|  | MIC USA300 | 0.88 |
|  | MIC Newman | 0.89 |
|  | Agglutination | 0.83 |
| Gradient boosting | MIC PA14 | 0.98 |
|  | MIC LESB58 | 1.00 |
|  | MIC USA300 | 0.99 |
|  | MIC Newman | 0.99 |
|  | Agglutination | 1.00 |

\*R<sup>2</sup>: for decision trees, as a single tree is fit, 20% of the sample datapoints were randomly selected for validation, so two R<sup>2</sup> values were obtained, one for training and one for validation. For RF and GB, it was possible to perform cross-validation (out-of-bag sampling with replacement) for each tree in the ensemble, so a single R<sup>2</sup> is reported for the ensemble. <sup>+</sup>: R<sup>2</sup> values of exactly 1 were obtained for both training and validation datasets.

**Table 5.** Fit metrics of the classification models.

| Method | Output variable | Precision* |  | Recall* |  | F1 score* |  |
| --- | --- | --- | --- | --- | --- | --- | --- |
| Decision tree | MIC PA14 | 1 | 1 | 1 | 1 | 1 | 1 |
|  | MIC LESB58 | 1 | 0.5 | 1 | 0.46 | 1 | 0.48 |
|  | MIC USA300 | 1 | 0.45 | 1 | 0.41 | 1 | 0.43 |
|  | MIC Newman | 1 | 0.44 | 1 | 0.36 | 1 | 0.4 |
|  | Agglutination | 1 | 0.61 | 1 | 0.58 | 1 | 0.58 |
| Random forest | MIC PA14 | 1 | 1 | 1 | 1 | 1 | 1 |
|  | MIC LESB58 | 1 | 1 | 1 | 1 | 1 | 1 |
|  | MIC USA300 | 1 | 0.46 | 1 | 0.5 | 1 | 0.48 |
|  | MIC Newman | 1 | 0.46 | 1 | 0.5 | 1 | 0.48 |
|  | Agglutination | 1 | 0.75 | 1 | 0.86 | 1 | 0.75 |
| Gradient boosting | MIC PA14 | 1 | 1 | 1 | 1 | 1 | 1 |
|  | MIC LESB58 | 1 | 0.5 | 1 | 0.38 | 1 | 0.43 |
|  | MIC USA300 | 1 | 0.45 | 1 | 0.45 | 1 | 0.45 |
|  | MIC Newman | 1 | 0.5 | 1 | 0.95 | 1 | 0.81 |
|  | Agglutination | 1 | 0.67 | 1 | 0.5 | 1 | 0.44 |

\*Precision, recall and F1 score are given as macro-average values (simple mean between the values for each class); the first value is for the training dataset, and the second, for the validation dataset. The precision, recall and F1 score for each class are shown in the Supporting information.

Classification models obtained good fits on the training dataset (achieving precision and recall of 1 for all methods and output variables) but failed to perform well on the testing dataset. Very few datapoints of MIC output variables (1 or 2) were available for class 1, which limited the fitting quality of the testing dataset (see Supporting information). Furthermore, values of 1 for precision, recall and F1 score of the testing dataset (Table 5) may give an incorrect sense of good fit 1 (see Supporting information). However, there was an almost even split of classes for hemagglutination (H_c_) output, although the fit of the testing dataset was marginally better than what was observed for MIC output variables for decision trees and RF models (i.e., a higher F1 score).

For these reasons, it was decided to continue with the modelling approach using regression models.

Regression models also achieved good fits (Table 4). For decision trees, the fits to both training and testing (validation) datasets were exactly 1, while for RF and GB, a single R^2^ value is used to represent the fit, as cross-validation was performed during fitting, and indicates good fits as well. Therefore, we plotted the actual value versus predicted value for regression RF and GB models (Figure 2) and the regression GB model has better predictability (Figure 2A).

**Figure 2.**
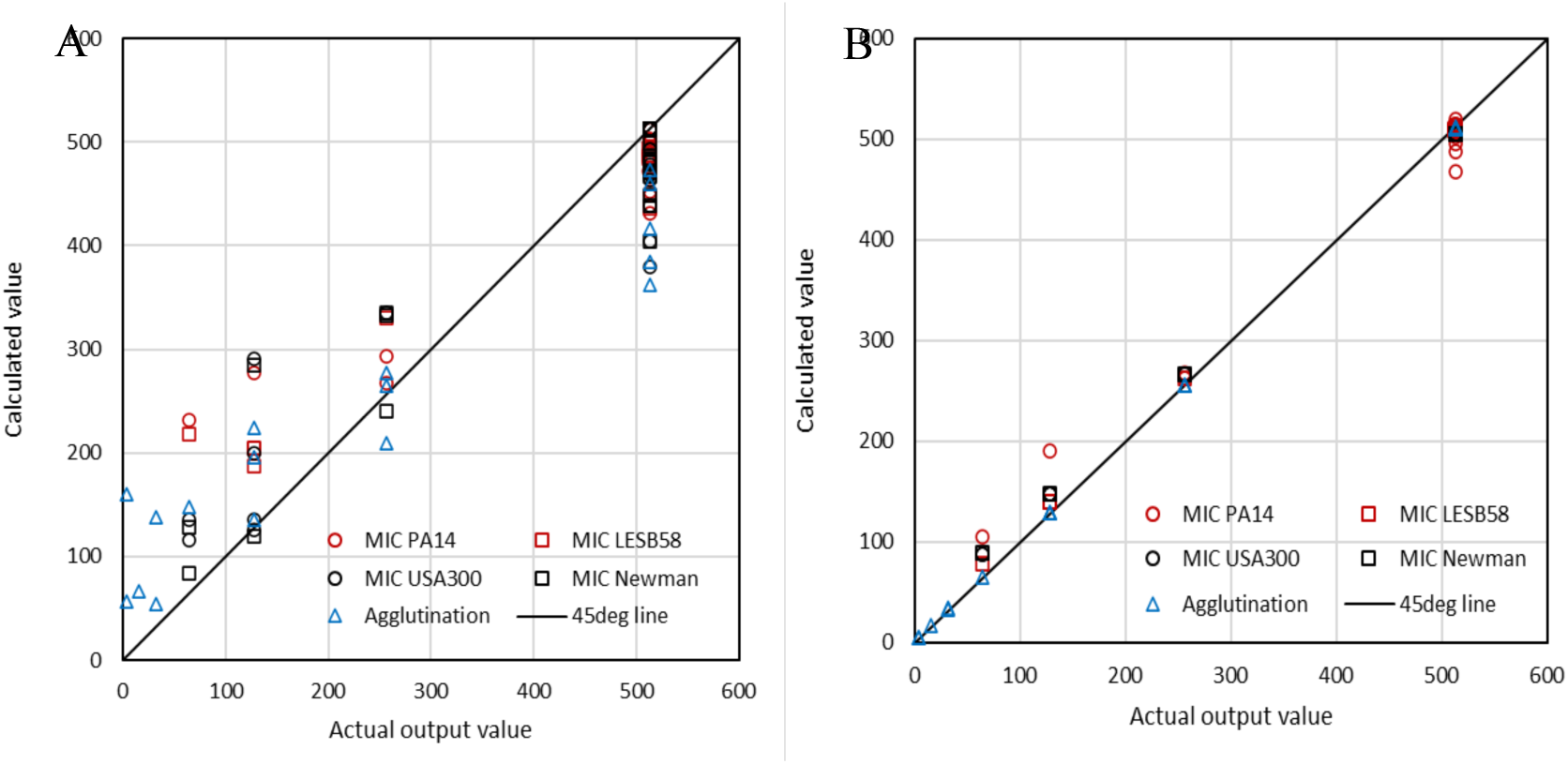
Predicted versus actual value for all output variables with A) RF regression model (left) and B) GB regression model (right).

In addition to considering the model fit, reproducibility of feature importance and consistency of generated tree structure were included in the analysis for selection of the best performing model(s).

### Reproducibility and consistency of the ML models

Reproducibility of feature importance between different models was tested by performing 5 runs in which distinct datapoints were selected for validation/cross-validation, and with separate hyperparameter optimisation (with 100 random trials). The average feature importance for each feature and each output variable is shown in Table 6 for regression models (the data for classification models is shown in the Supporting information). The optimised hyperparameter values, fitting metrics and individual feature importance values for each run can be seen in Supporting information.

**Table 6.**
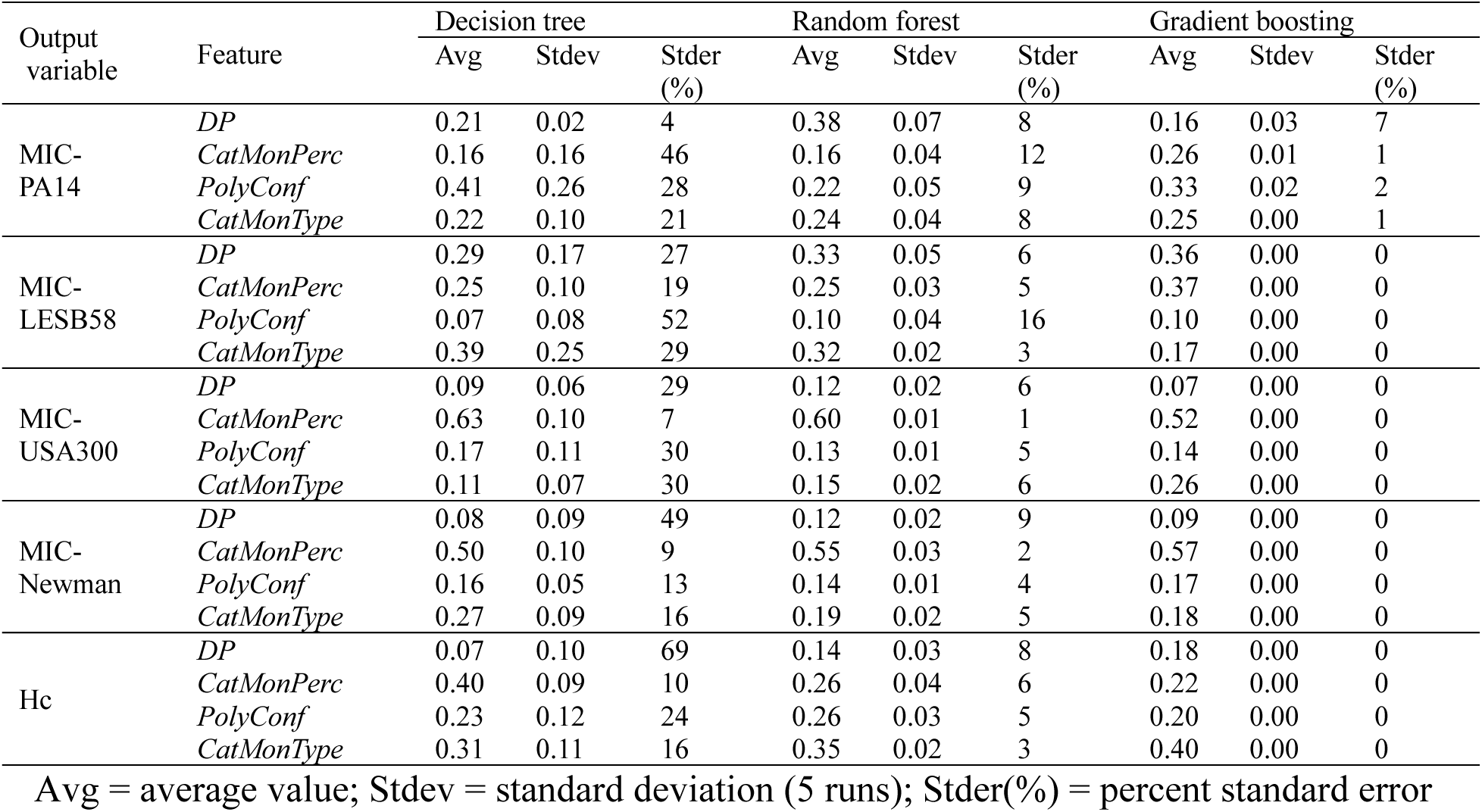
Average feature importance, standard deviation and error for regression models.

For regression models, the decision tree had the highest standard error between all runs, averaging 20-34% for the different output values. RF and GB had average standard errors ranging between 4-9% and 0-3%, respectively. Classification models (Table S4 in the Supporting information) exhibited larger standard errors: 28-51% for decision tree, 17-37% for RF, and 23-64% for GB.

Regression RF and GB models were the most reproducible, and this can be more easily visualised by plotting the feature importance of different runs in spider plots (Figure 3). The plots for all other regression and classification models are in the Supporting information (Figures S2 to S6).

**Figure 3.**
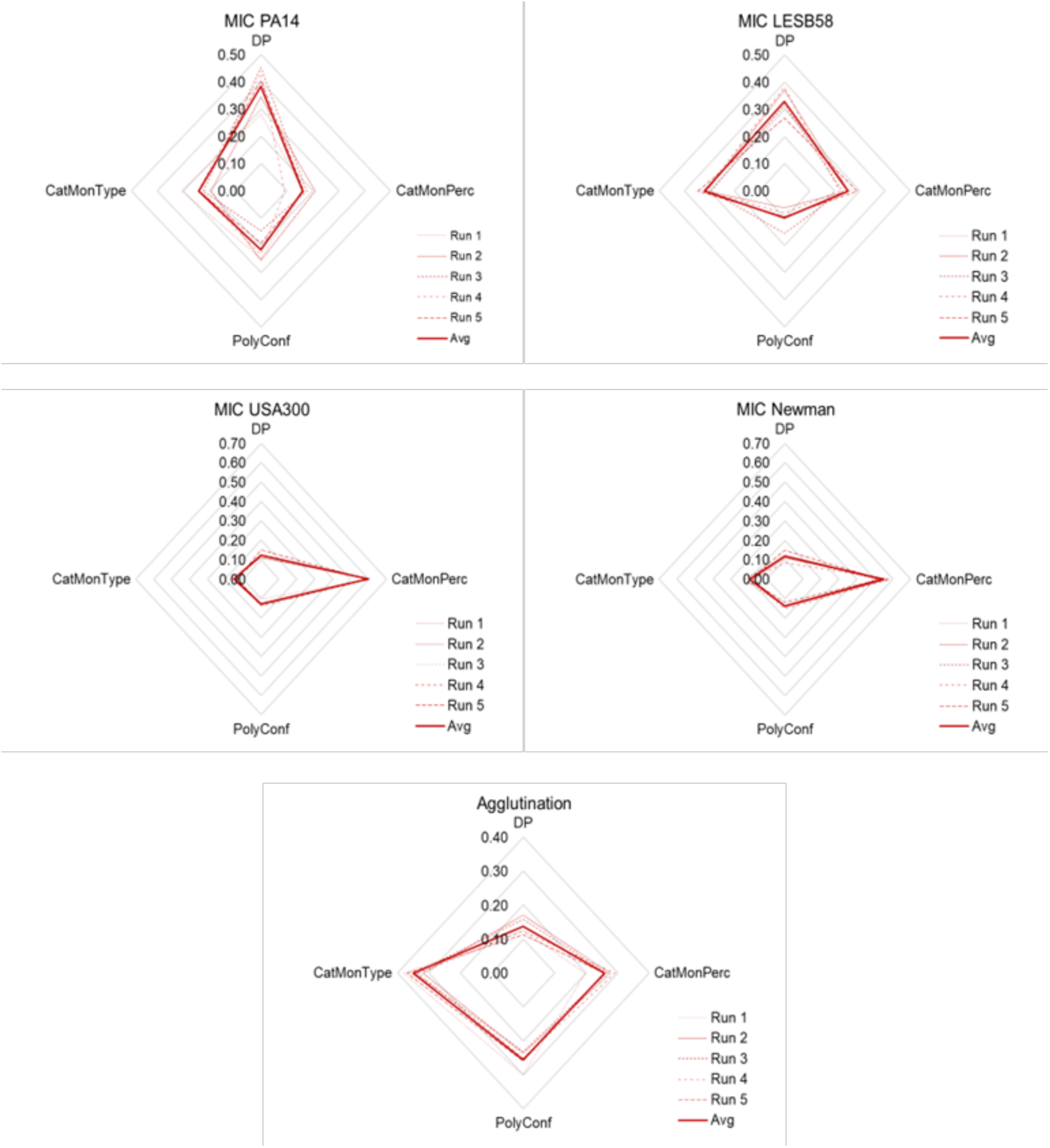
Reproducibility test of feature importance for regression RF models. For each output variable (MIC PA14, MIC LESB58, MIC USA300, MIC Newman, and Agglutination), five different runs were performed, and the importance of each feature (degree of polymerisation [DP], type of cationic monomer [CatMonType], percentage of cationic monomer [CatMonPerc], and polymer confirmation [PolyConf]) calculated for each run. The average across all runs is also shown. In tree-based models, feature importance is computed during model fitting and derives from node impurity calculations.

The consistency of tree structure between different runs was also analysed. For RF models, tree structures are individually less impactful given that the average of all trees’ predictions is taken into account during bagging. This analysis is discussed in detail in the supporting information. Briefly, the decision tree models mostly fit distinct tree structures across different runs, and this resulted in the high variability of average feature importance. For GB regression models, while the initial trees in the sequence have presented slight structural changes between runs, the final trees were either identical or had node splits at deeper levels (which have a smaller impact on feature importance).

In general, the classification models had low fitting scores at the testing stage of the dataset and with inability to reproduce the average feature importance distributions after multiple runs. On the other hand, RF and GB regression models fit the data well and were reproducible across distinct runs. This indicates that these two models best capture the input-output relationships of the data and were therefore applied to further ML data analysis.

### Feature importance and average SHAP values

Based on both model fit, feature importance reproducibility and consistency of tree structures, as discussed previously, the regression RF and GB models were selected for representing the dataset and further SHAP analysis.

### Feature importance for RF and GB regression models

From the average feature importance values (Table 6, Figure 3), both RF and GB models predicted similar feature importance distributions for MIC USA300 and MIC Newman, with the percentage of the cationic monomer being the greatest contributing input (feature) with 50-60% contribution, followed by the type of cationic monomer and polymer conformation, and finally the degree of polymerisation, with less than 20% contribution.

MIC for *P. aeruginosa* strains exhibited variable feature contributions, for both the models and the strains. Based on the RF regression model, the order of features contributions for MIC PA14 was DP > type of cationic monomer > polymer conformation and cationic monomer ratio. While for MIC LESB58, the order was similar but the DP and type of cationic monomer were the most contributing features closely at around 30%, then the cationic ratio and polymer conformation with about 10% contribution.

With the GB regression model, for MIC PA14, more importance was placed on polymer conformation (33% versus 22%) and percentage of cationic monomer (26% versus 16%), rather than on DP (16% versus 38%), relative to the RF model. For MIC LESB58, GB model placed more importance on the percentage of cationic monomer (37% versus 25%), but the importance was relatively the same for DP and polymer conformation. The GB model also placed less importance on the type of cationic monomer (17% versus 32%) than RF.

For agglutination, RF and GB placed similar levels of importance on the same features. The type of cationic monomer was the most important feature (35-40%), followed by polymer conformation (22-26%) and cationic monomer ratio (26-20%).

### Comparing feature importance and average SHAP values for RF and GB regression models

SHAP values explain how each feature impacts the expected output values in a dataset. While feature importance only provides the magnitude of the contribution for the whole dataset, SHAP provides both the magnitude and the type of contribution (positive or negative) for each datapoint in the dataset. The advantage of feature importance is that it is calculated as part of the tree fitting procedure (refer to the equations presented earlier in the Experimental section), and no extra computational work is needed to obtain those values. SHAP values were averaged over the dataset and compared to the feature importance (Figure 4). The values in the graphs are listed in the supporting information (Tables S17 and S18).

**Figure 4.**
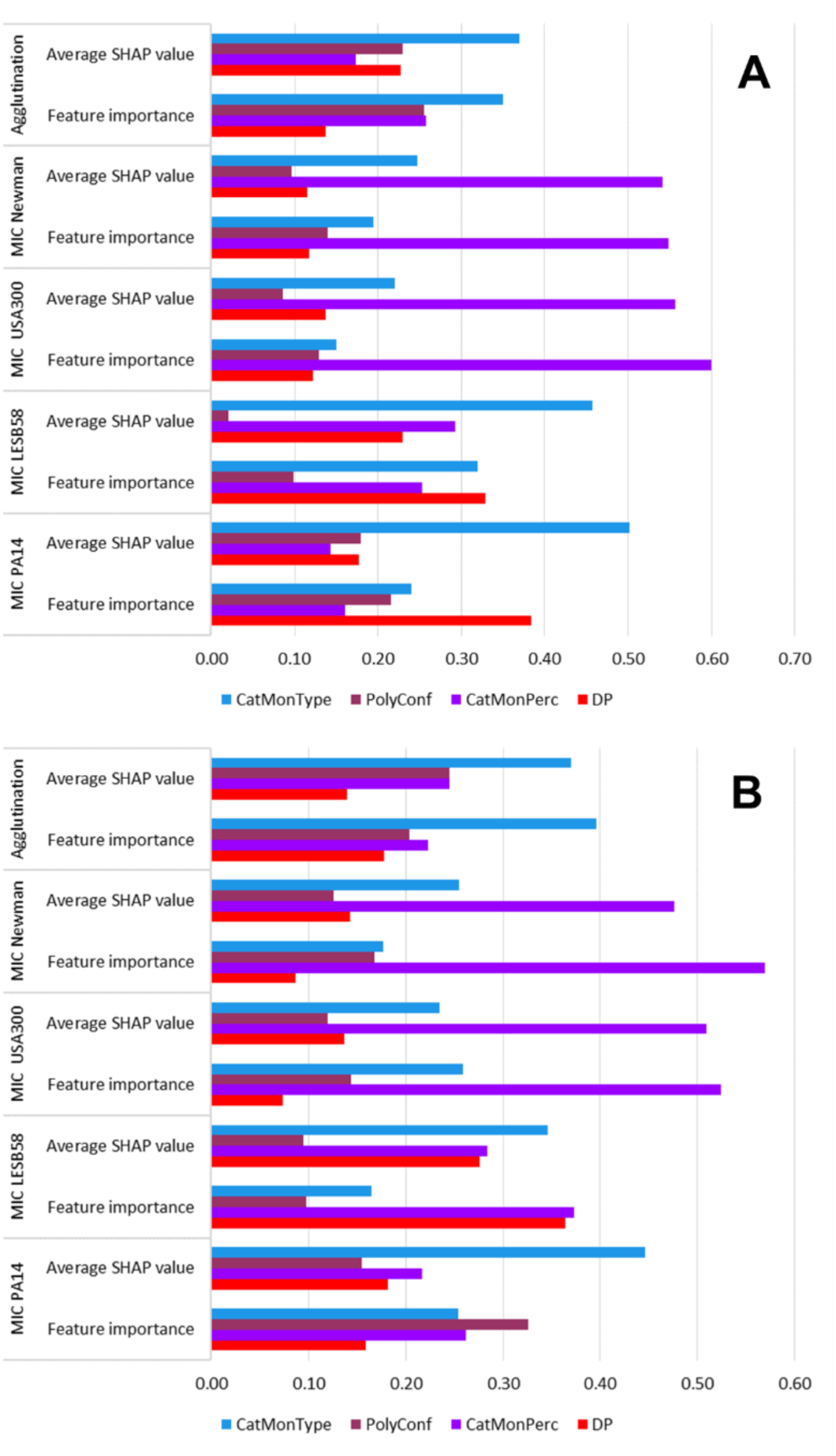
Comparison of feature importance and average SHAP values for all output variables (MIC PA14, MIC LESB58, MIC USA300, MIC Newman, and Agglutination). The importance and SHAP value of each model feature (input variable) - degree of polymerisation (DP), type of cationic monomer (CatMonType), percentage of cationic monomer (CatMonPerc), and polymer confirmation (PolyConf) – is shown on a scale of 0 to 1, and these values add up to 1. This comparison is illustrated for RF regression (A) and GB regression models (B).

In the RF model, the average SHAP value places less importance on DP and more importance on the type of cationic monomer than feature importance, for both MIC PA14 and MIC LESB58. The importance of polymer conformation and percent of cationic monomer remained similar for MIC PA14, while for MIC LESB58, the importance of polymer conformation was minimal with the average SHAP value. For the other output variables (MIC USA300, MIC Newman and agglutination), feature importance and average SHAP values were equivalent, and there was not much variation in their distributions.

With the regression GB model, SHAP values were also similar to feature importance distributions for MIC USA300, MIC Newman and agglutination. For MIC PA14, the average SHAP value allocated greater importance to the type of cationic monomer and reduced importance to polymer conformation relative to average feature importance. For MIC LESB58, average SHAP also placed more importance on the type of cationic monomer, but less importance to the degree of polymerisation and percentage of cationic monomer.

Comparing the average SHAP values from the RF and GB models, it should be noted that their distributions are more similar than those of feature importance from the same RF and GB models. The average SHAP values from different models give more consistent results for the overall dataset than feature importance.

### Datapoint-specific SHAP values

SHAP values for each data point represent the marginal contribution of each feature relative to the average output value for the whole dataset. In other words, SHAP value gives how much each feature contributes to increase or decrease a data point’s output compared to the average output value. We visualised the SHAP values via waterfall curves and beeswarm plots.

In the beeswarm plots (Figure 5), each dot on the plots represents one data point. The x-axis is the SHAP value. A value of zero means the data point does not contribute to changing the output value. A negative value means that the feature decreases the value of the output while a positive value means it increases it, all in relation to the average value of the variable. The average values between the two models (Table 7) are slightly different, as each model provides a different estimate for each datapoint. The colours of the dots are related to the feature value with the colour bar on the right. For features that are inherently categorical but treated as continuous during modelling, the high-low grading can be related to the original feature class by referring to Table 1 and Table 2. For example, low CatMonType corresponds to cationic monomer 1 (AEAM), and medium-high PolyConf to polymer conformation 3 (triblock copolymer).

**Figure 5.**
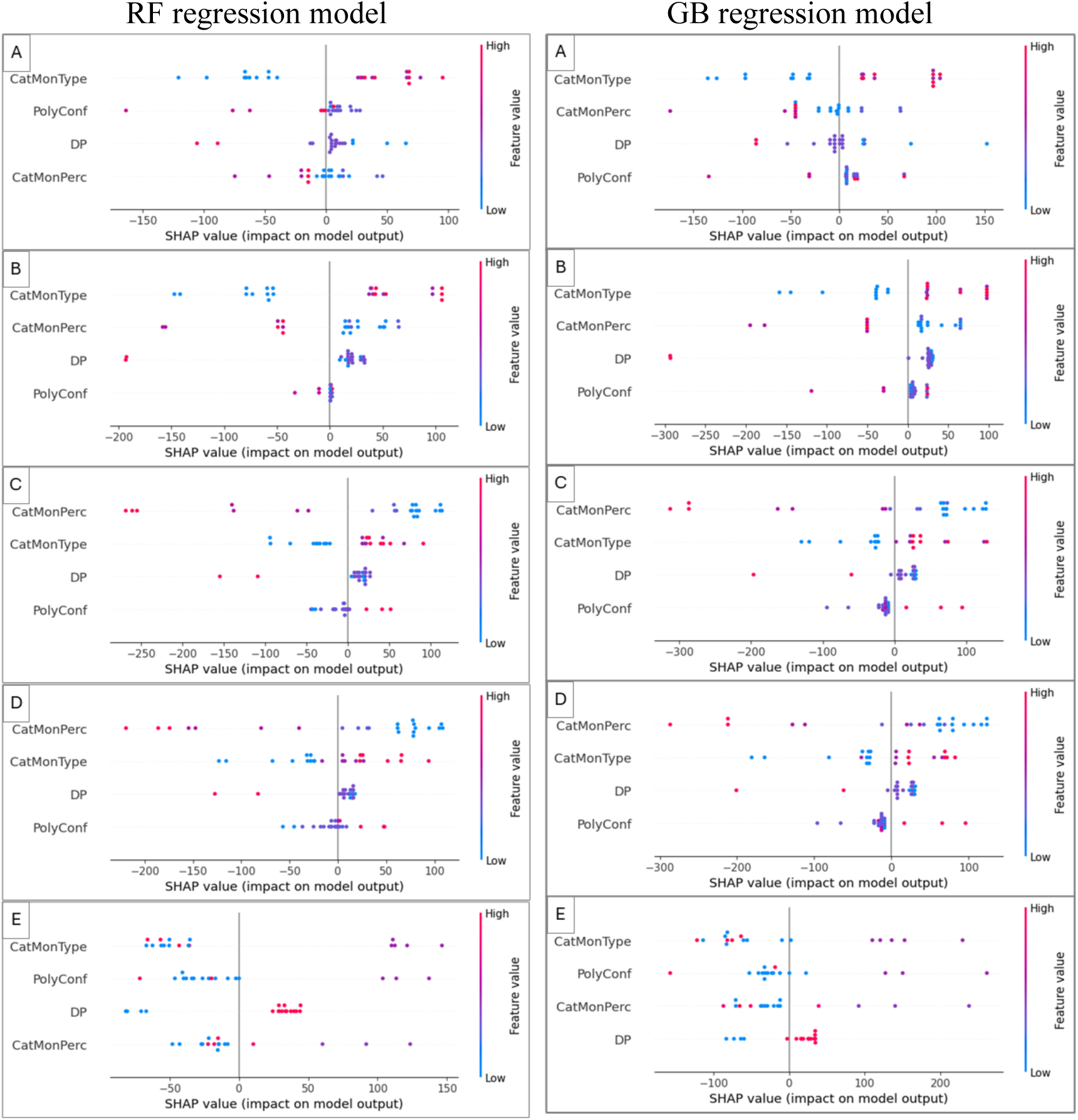
Beeswarm plots for RF regression models (left) and GB regression model (right). The colour bar gives the magnitude of continuous features and indicates the class for categorical features. A: MIC PA14; B: MIC LESB58; C: MIC USA300; D: MIC Newman; E: Hemagglutination.

**Table 7.**
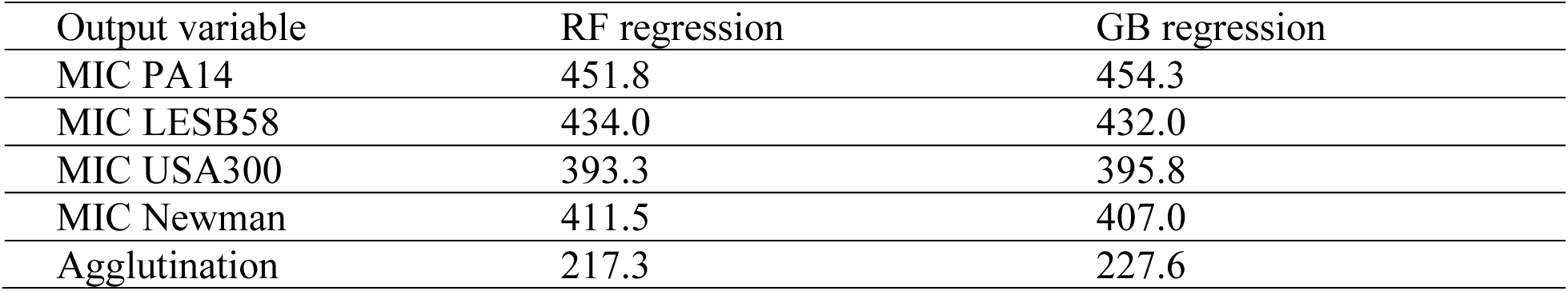
Dataset-averaged output values, E[f(x)], in µg/mL for RF and GB regression models.

| Output variable | RF regression | GB regression |
| --- | --- | --- |
| MIC PA14 | 451.8 | 454.3 |
| MIC LESB58 | 434.0 | 432.0 |
| MIC USA300 | 393.3 | 395.8 |
| MIC Newman | 411.5 | 407.0 |
| Agglutination | 217.3 | 227.6 |

Therefore, the beeswarm plots can be used to identify feature values which contribute to moving the output closer to the desired value. For all four MIC outputs, a small output value is desirable, as this means a greater potency. On the contrary, higher output values are desirable for hemagglutination (C_H_).

Comparing the beeswarm plots for RF and GB models (Figure 5), the SHAP value spread is similar for all outputs. While there are some differences in the magnitude of the SHAP values, the colour distribution of the dots remains the same for each output variable. For example, focusing on each of the four features for MIC PA14 and specifically on the negative side of the x-axis (Figure 5A), it can be seen that the feature values leading to negative SHAP are: low for type of cationic monomer, high for DP, medium-high for polymer conformation and medium-high to high for cationic monomer ratio (although some low CatMonPerc values are also slightly negative). According to this, DP 100 triblock copolymer, with over 50% AEAM will result in the lowest MIC-PA14, for example. A similar analysis was performed for the other outputs and the results are summarised in Table 8.

**Table 8.**
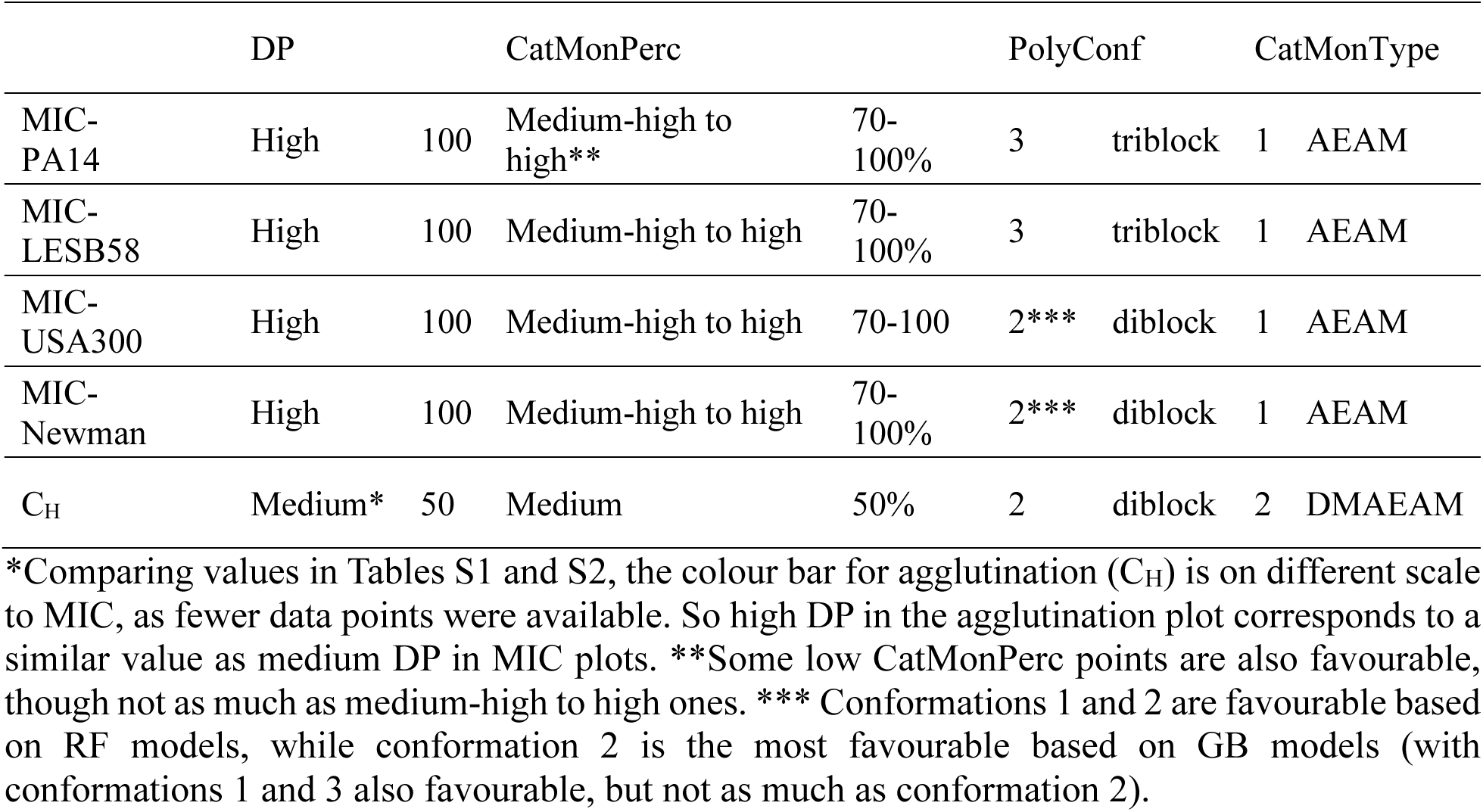
Feature values that contribute to moving the output closer to its desired value.

It is worth mentioning that the most visible difference in SHAP value spread occurs for the polymer conformation feature in MIC USA300 and MIC Newman models (Figure 5C&D). While for RF models, homopolymer and diblock (polymer conformations 1 and 2 in blue and light purple dots) are more favourable (i.e., more negative), for GB models, diblock (polymer conformation 2) has the most negative SHAP value, thus the most favourable. Conformations 1 (as in RF) and 3 (homopolymer and triblock) also have negative SHAP values and are favourable.

Caution should be taken when interpreting beeswarm plots, as the features are considered separately, so any potential correlations between features could have been overlooked. These potential correlations may be observed when looking into the combined effect of the SHAP values of all features on specific datapoints. For this purpose, waterfall plots are used.

Waterfall plots (Figure 6, Figure 7 and Figure 8) are datapoint-specific and show how each feature contributes (both in magnitude and direction) to taking the output value from the sample-averaged value (E[f(x)]) to the predicted output value (f(x)) for the selected datapoint. Here, we will discuss the waterfall plots for the best-performing datapoints in each output variable (MIC ≤ 64 µg/mL and agglutination concentration > 256 µg/mL).

**Figure 6.**
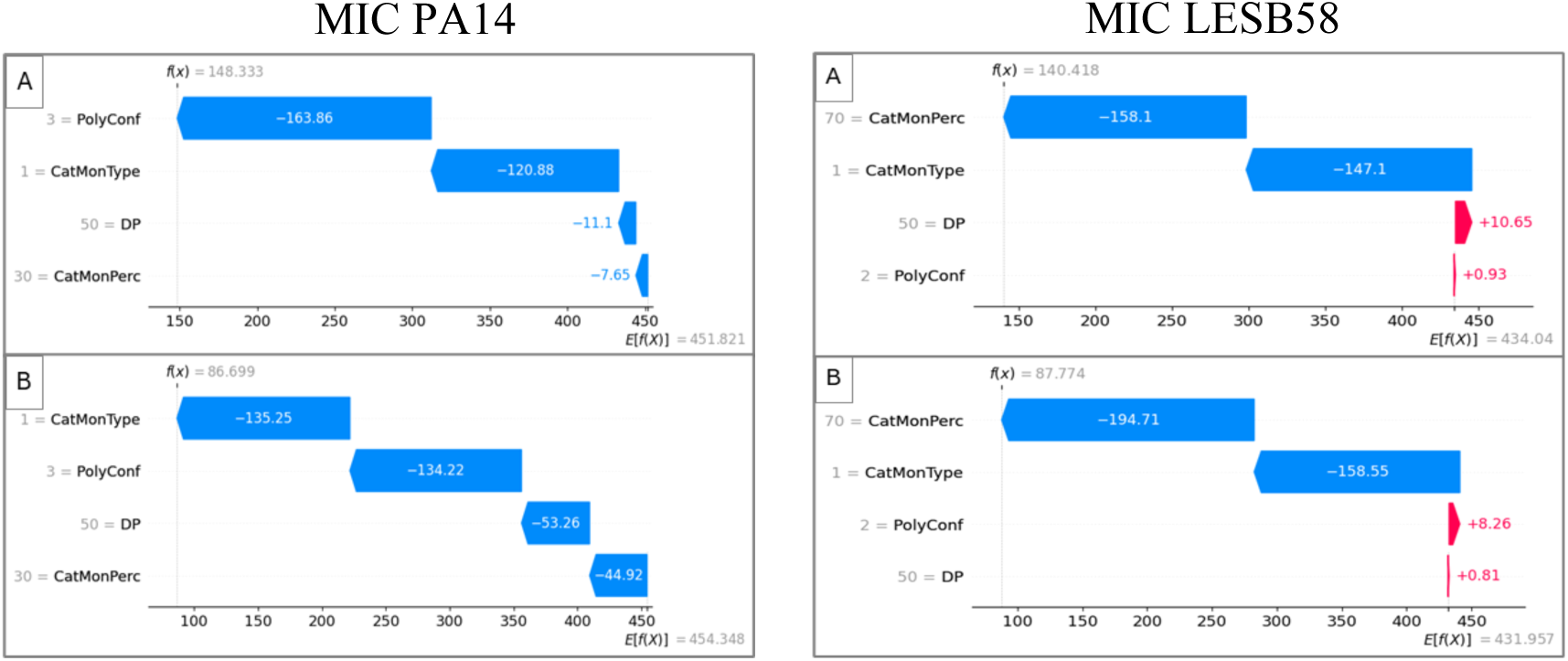
Waterfall plots for best-performing MIC PA14 (data point 15) and MIC LESB58 (data point 8) SAMP designs for each model: A (RF model) and B (GB model). Waterfall plots depict datapoint-specific SHAP values. Reading the plots starts at the dataset-averaged output value (E[fx)], as listed in Table 7). Then each feature’s SHAP value (identified by red arrows if positive, or blue arrows if negative) adds to or subtracts from the dataset-averaged output value, giving as the final answer the datapoint-specific output value (f(x)) for a given set of feature values.

**Figure 7.**
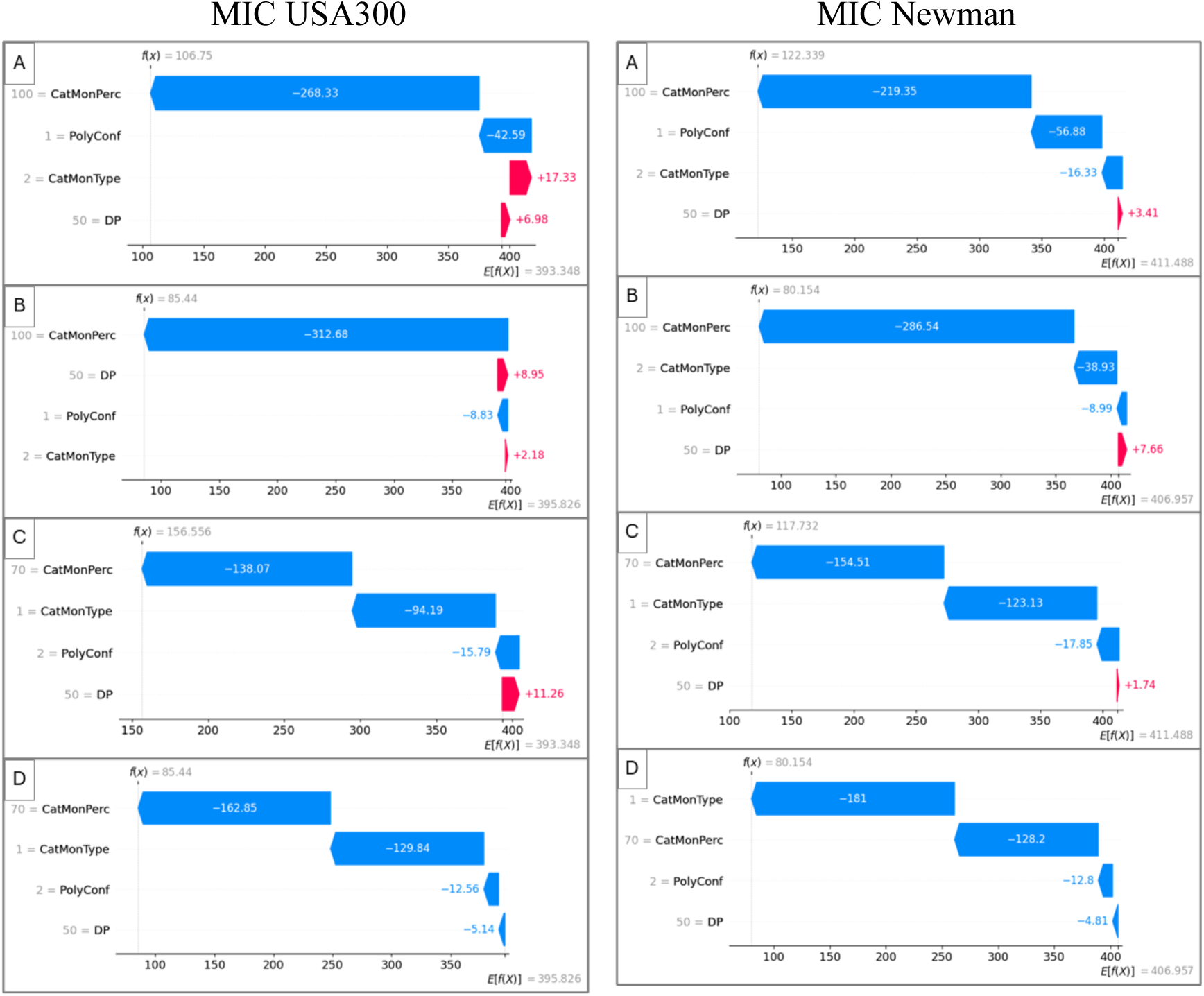
Waterfall plot for best-performing MIC USA300 (left) and MIC Newman (right) SAMP designs (datapoints 5 and 8). A: RF model datapoint-specific SHAP values for datapoint 5; B: GB model datapoint-specific SHAP values for datapoint 5; C: RF model datapoint-specific SHAP values for datapoint 8; D: GB model datapoint-specific SHAP values for datapoint 8. Waterfall plots depict datapoint-specific SHAP values. Reading the plots starts at the dataset-averaged output value (E[fx)], as listed in Table 7). Then each feature’s SHAP value (identified by red arrows if positive, or blue arrows if negative) adds to or subtracts from the dataset-averaged output value, giving as the final answer the datapoint-specific output value (f(x)) for a given set of feature values.

**Figure 8.**
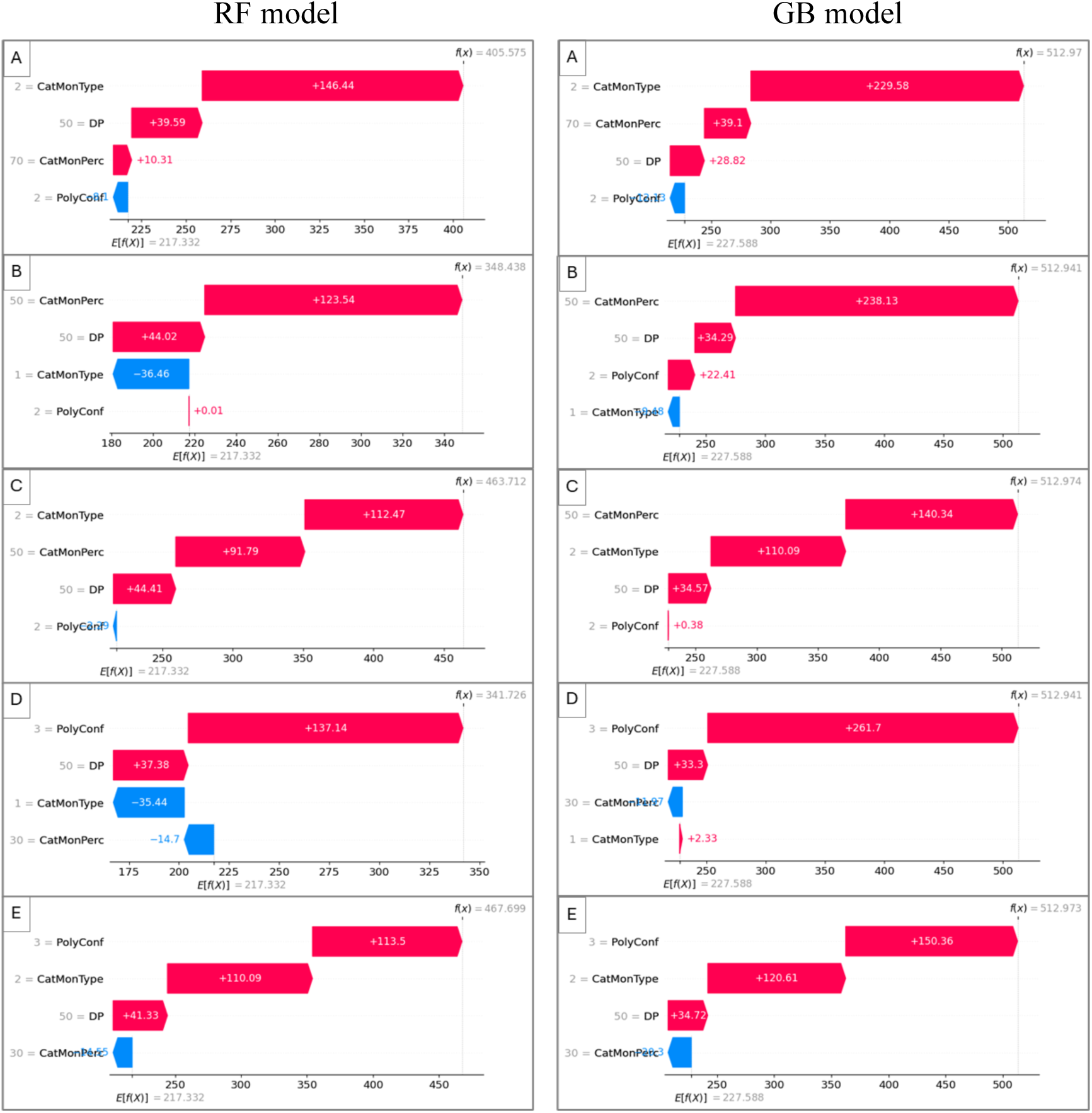
Waterfall plot for best-performing Agglutination SAMP designs using the RF model (left) and GB model (right). A: datapoint 7’; B: datapoint 8’; C: datapoint 10’; D: datapoint 11’; E: datapoint 14’. Waterfall plots depict datapoint-specific SHAP values. Reading the plots starts at the dataset-averaged output value (E[fx)], as listed in Table 7). Then each feature’s SHAP value (identified by red arrows if positive, or blue arrows if negative) adds to or subtracts from the dataset-averaged output value, giving as the final answer the datapoint-specific output value (f(x)) for a given set of feature values.

The waterfall plots for the best-performing MIC PA14 datapoint obtained from both the RF and GB models are shown in (Figure 6). For both models, the datapoint-specific SHAP values suggest that polymer conformation 3 (triblock) and type of cationic monomer 1 (AEAM) are the largest contributors to reducing the MIC value. The GB model places more contribution on the other two features, leading to a predicted value (86.7 µg/ml) closer to the actual MIC (64 µg/ml).

Interestingly, while based on the beeswarm plots (Figure 5), high DP was predicted to be beneficial, the best-performing datapoint in waterfall plot was a medium DP value (DP50). This illustrates how combined with polymer conformation 3 (triblock) and cationic monomer 1 (AEAM), a lower DP can be favourable. Although it should be noted that a datapoint with a higher DP and these same conformation and cationic monomer values was not included. This highlights the importance of investigating the relationships between features to further inform, confirm, or challenge insights gained from a beeswarm plot.

Similarly to MIC PA14, MIC LESB58 only had one best-performing datapoint, datapoint 8 (Figure 6). The type of cationic monomer was also AEAM (1), but differently from MIC PA14, LESB58 had the most beneficial contribution from the percent of cationic monomer, at a medium-high value (70%).

Two datapoints performed well for MIC USA300 (datapoints 5 and 8). For both datapoints and in both models (Figure 7), cationic monomer percent was the most favourable feature, at a medium to high value (70-100%). Homopolymers and diblock copolymers were both favourable (classes 1 and 2). AEAM monomer was the second-best contributor. Both models explained the datapoint performance in similar ways, with only a small difference in the DP SHAP value for data point 8 (a slightly positive value for RF, but slightly negative for GB). MIC Newman followed similar trends (Figure 7), except that for datapoints 5 and 8. For the GB model datapoint 5, the type of cationic monomer was the second most contributing feature instead of polymer conformation. For datapoint 8, the type of cationic monomer was contributing more than the percent of cationic monomer. However, both models and datapoints considered the percentage of cationic monomer at medium-high amounts (70%) and cationic monomer AEAM as the two most contributing features, similarly to MIC USA300.

On the other hand, hemagglutination had 5 datapoints that performed well (datapoints 7’, 8’, 10’, 11’, and 14’). Different features take on different levels of importance depending on the datapoint, but some trends can be observed overall (Figure 8). Firstly, cationic monomer percents of at least 50% are beneficial. Triblock copolymer conformation and DMAEAM monomer, either combined or alone, contributed in a favourable way. DP of 50 was also favourable. Both models agree on these observations. It is important to compare these with the results expected from the beeswarm plot, which identified DMAEAM as the best monomer but diblock as the best polymer conformation (Table 8). But from the waterfall plot (Figure 8), diblock copolymers were slightly unfavourable (or had no effect) for the best-performing datapoints 7’, 8’ and 10’.

It should be noted that the most contributing features for the best-performing datapoints do not always match in type and amount of contribution with the most contributing features from the average SHAP value or feature importance contribution distributions (Figure 4). The average SHAP value and feature importance give dataset-averaged contributions, and so they also take into account datapoints that did not perform as well. Hence it may be possible that some of the most effective features to the output, contributed in a negative way.

## Discussion

Amongst the models tested, the tree ensemble regression methods had the best fit and reproducibility after multiple runs, which was different to a similar study by Kundi *et al.* that proposed the use of decision tree classification model with another set of descriptors.^64^ Here, after running FI and SHAP analysis for GB and RF regression models, the features importance and their values that provided the best performance were elucidated for each output as follows (in order of higher to lower contribution): For MIC-*P. aeruginosa* (both strains): type of cationic monomer (AEAM), the cationic ratio (70-100%), chain length (DP100), and polymer conformation (triblock). For MIC-*S. aureus* (both strains): the cationic ratio (70-100%), type of cationic monomer (AEAM), chain length (DP100), and polymer conformation (diblock). For Hemagglutination: type of cationic monomer (DMAEAM), polymer conformation (diblock), chain length (DP100), and the cationic ratio (50%).

There are similarities between structure-activity relationship patterns detected here to our SAR analysis in previous work.^44^ Beeswarm plot model feature suggestions align with some of the experimental observation of the best performing SAMPs design features, especially for polymer conformation, type and percentage of cationic monomer. However, complete alignment is unexpected, firstly because beeswarm plot analysis does not consider any potential feature correlations. Secondly, insights based on the models may contain feature combinations which have not been experimentally collected, thus not allowing for a direct match, but providing additional information. According to beeswarm, the cationic moiety and the cationic ratio are the most contributing features to the potency against *S. aureus* strains, which matches our conclusion based on experimental observation. Interestingly, feature importance and average SHAP value analysis suggests that DP is the least important feature for activity against *S. aureus* strains, which corroborates the idea that all other features being at their optimal values, the DP value can vary in the 50-100 range without affecting performance.

This shows the suitability of the descriptors used here as they were all contributing to the activity and selectivity of the polymers. The SHAP analysis can be used as guideline to design future SAMPs libraries of similar structures with higher success rate. For example, utilising the cationic monomer AEAM in high ratio (>50%) contribute to high potency and can be used in segmented conformation (diblock or triblock) with DP100. It must be noted that the dataset used here is limited and can benefit from more datapoints that cover a wider range of values.

On the other hand, a different model can generate completely different patterns. Kundi *et al.* work with other descriptors and model recommended alternative design. SHAP values in their work recommended increasing NIPAM ratio over 0.4, maintaining the clogP between 0.5 and -2, and even the omission of AEAM to increase the likelihood of potent polymers.^64^ Hence, it is important to evaluate different models in small, controlled library for the most suitable model to be used with larger dataset, and to identify the relevant input values to be used that suit the purpose.

## Conclusion

Screening through the library of 23 potential antimicrobial polyacrylamides, their performance was impacted by the key features: type of cationic monomer, cationic ratio, DP and polymer architecture. These features were useful parameters in the machine learning models and can be utilised to predict future candidates. Among the models operated in this pilot study, tree-ensemble regression models; GB and RF, were good fit of the dataset and produced consistent feature importance values across multiple runs, hence, have great potentials forecasting prospective SAMPs candidates. SHAP and features importance outcomes highlighted the importance of the type and number of cationic moieties on the polymers’ activity. Although the machine learning predictions in this work were consistent within this dataset, it still requires manifold data points to produce informed predictions. This can be achieved by feeding the model presented in this work (regression gradient boosting) with the published libraries of SAMPs using the proposed features as descriptors.

## Acknowledgements

LD thanks Cara for the provision of a fellowship.

